# *Sox4* drives *Atoh1-*independent intestinal secretory differentiation toward tuft and enteroendocrine fates

**DOI:** 10.1101/183400

**Authors:** Adam D Gracz, Leigh Ann Samsa, Matthew J Fordham, Danny C Trotier, Bailey Zwarycz, Yuan-Hung Lo, Katherine Bao, Joshua Starmer, Noah F Shroyer, R. Lee Reinhardt, Scott T Magness

## Abstract

**Background & Aims:** The intestinal epithelium is maintained by intestinal stem cells (ISCs), which produce post-mitotic absorptive and secretory epithelial cells. Initial fate specification toward enteroendocrine, goblet, and Paneth cell lineages is dependent on *Atoh1*, a master regulator of secretory differentiation. However, the origin of tuft cells, which participate in Type II immune responses to parasitic infection, is less clear and appears to occur in an *Atoh1*-independent manner. Here we examine the role of *Sox4* in ISC proliferation and differentiation.

**Methods:** We used mice with intestinal epithelial-specific conditional knockout of *Sox4* (Sox4^fl/fl^:vilCre; Sox4cKO) to study the role of *Sox4* in the small intestine. Crypt- and single cell-derived organoids were used to assay proliferation and ISC potency between control and Sox4cKO mice. Lineage allocation and genetic consequences of *Sox4* ablation were studied by immunofluorescence, RT-qPCR, and RNA-seq. *In vivo* infection with helminths and *in vitro* cytokine treatment in primary intestinal organoids were used to assess tuft cell hyperplasia in control and Sox4cKO samples. *Atoh1*^*GFP*^ reporter mice and single cell RNA-seq (scRNA-seq) were used to determine co-localization of SOX4 and *Atoh1*. Wild-type and inducible *Atoh1* knockout (Atoh1^fl/fl^:vilCreER; Atoh1iKO) organoids carrying an inducible *Sox4* overexpression vector (Sox4OE) were used to determine the role of *Atoh1* in *Sox4* driven secretory differentiation.

**Results:** Loss of *Sox4* impairs ISC function and secretory differentiation, resulting in decreased numbers of enteroendocrine and tuft cells. In wild-type mice, SOX4+ cells are significantly upregulated following helminth infection coincident with tuft cell hyperplasia. *Sox4* is activated by IL13 *in vitro* and Sox4cKO knockout mice demonstrate impaired tuft cell hyperplasia and parasite clearance following infection with helminths. A subset of *Sox4*-expressing cells colocalize with *Atoh1* and enteroendocrine markers by scRNA-seq, while *Sox4+*/*Atoh1*-cells correlate strongly with tuft cell populations. Gain-of-function studies in primary organoids demonstrate that *Sox4* is sufficient to drive both enteroendocrine and tuft cell differentiation, and can do so in the absence of *Atoh1*.

**Conclusion:** Our data demonstrate that *Sox4* promotes enteroendocrine and tuft cell lineage allocation independently of *Atoh1*. These results challenge long-standing views of *Atoh1* as the sole regulator of secretory differentiation in the intestine and are relevant for understanding host epithelial responses to parasitic infection.

## Introduction

The intestinal epithelium is essential for both digestive and barrier function and contains a diverse complement of post-mitotic cells which carry out complex physiological functions critical for homeostasis. Functional cell types in the intestine can be broadly subdivided into: (a) absorptive enterocytes and (b) secretory lineages, which include antimicrobial-producing Paneth cells, mucus-producing goblet cells, and hormone-secreting enteroendocrine cells (EECs) ^1^. Tuft cells, which represent a less well-characterized lineage, initiate immune responses to parasitic infections in the intestine ^2-5^.With the exception of long-lived Paneth cells, these post-mitotic lineages are subject to the rapid (5-7 day) and continuous turnover of the intestinal epithelium. Therefore, their numbers must be maintained through constant proliferation and differentiation of the intestinal stem cells (ISCs), which reside in a specialized stem cell niche at the base of the intestinal crypts.

Extrinsic signals from the niche induce regulatory programs in ISCs and their immediate progeny, transit-amplifying progenitor cells (TAs), to drive commitment to different cellular lineages. For example, loss of Notch signaling in ISCs and early progenitors is associated with the induction of secretory differentiation through the derepression of *Atoh1*, which is widely regarded as the master regulator of secretory fate ^6, 7^. Certain pathological settings can shift lineage allocation in the intestine, as exemplified by goblet and tuft cell hyperplasia in the setting of type II immune responses.

While tuft cells are also derived from ISCs, the transcriptional regulatory mechanisms governing their differentiation and hyperplastic response to infection remain less well characterized ^3^. This is due in part to evidence that tuft cells differentiate independently of *Atoh1*, but still appear to require downregulation of Notch ^6, 8, 9^. Recent studies have demonstrated upregulation of tuft cell numbers following *Atoh1* deletion *in vivo*, and systems biology approaches have supported an *Atoh1*-independent differentiation program for this lineage ^10^. Together, these data suggest an apparently paradoxical Notch-repressed, *Atoh1*-independepent mechanism for tuft cell specification.

Sry-box containing (Sox) factors are transcription factors with broad regulatory roles in stem cell maintenance and differentiation in many adult tissues, including the intestinal epithelium ^11, 12^. *Sox9* is known to contribute to Paneth cell differentiation, and previous work from our group has demonstrated that distinct expression levels of *Sox9* mark stem and progenitor cell populations from intestinal crypts, including label-retaining cells (LRCs) ^13-16^. The role of other Sox factors in ISC function and differentiation remain uncharacterized.

*Sox4* has been associated with the ISC genomic signature and is correlated with increased invasion/metastasis and decreased survival in colon cancer, but its role in intestinal epithelial homeostasis is unknown ^17, 18^. In the hematopoietic system, as well as the developing nervous system, pancreas, kidney, and heart, *Sox4* is required for both proper cell lineage allocation and maintenance of progenitor pools ^19-21^. The diverse regulatory potential of *Sox4*, coupled with its known role in a broad range of tissue-specific progenitor populations, led us to hypothesize that it may function as a regulator of ISC maintenance and differentiation. In this study we establish a role for *Sox4* in ISC homeostasis and regulation of tuft cell hyperplasia during parasitic helminth infection, which represents a novel pathway for tuft cell specification with important implications for human responses to parasitic disease.

## Results

### SOX4 is expressed at distinct levels in the ISC zone

To characterize *Sox4* expression, we first used *in situ* hybridization to detect *Sox4* mRNA. Consistent with previous reports, *Sox4* is restricted to the base of the intestinal crypts in the stem cell zone (Figure 1A) ^18^. Since the cellular resolution of *in situ* hybridization is low, we leveraged a previously characterized *Sox9*^*EGFP*^ reporter mouse to examine the expression of *Sox4* in different crypt cell types. Distinct *Sox9*^*EGFP*^ levels differentially mark active ISCs (*Sox9*^*low*^), secretory progenitors/reserve ISCs (*Sox9*^*high*^), TA progenitors (*Sox9*^*sublow*^), Paneth cells (*Sox9*^*EGFP-neg*^:CD24^+^), and non-Paneth post-mitotic cells (*Sox9*^*EGFP-neg*^:CD24^neg^) ^13, 14, 22, 23^. *Sox9*^*EGFP*^ populations were isolated by fluorescence-activated cell sorting (FACS) and analyzed by RT-qPCR. *Sox4* was upregulated in two populations: *Sox9*^*low*^, which is consistent with active ISCs, and *Sox9*^*high*^, which is consistent with secretory progenitors (Figure 1B) ^23, 24^.

**Figure 1.**
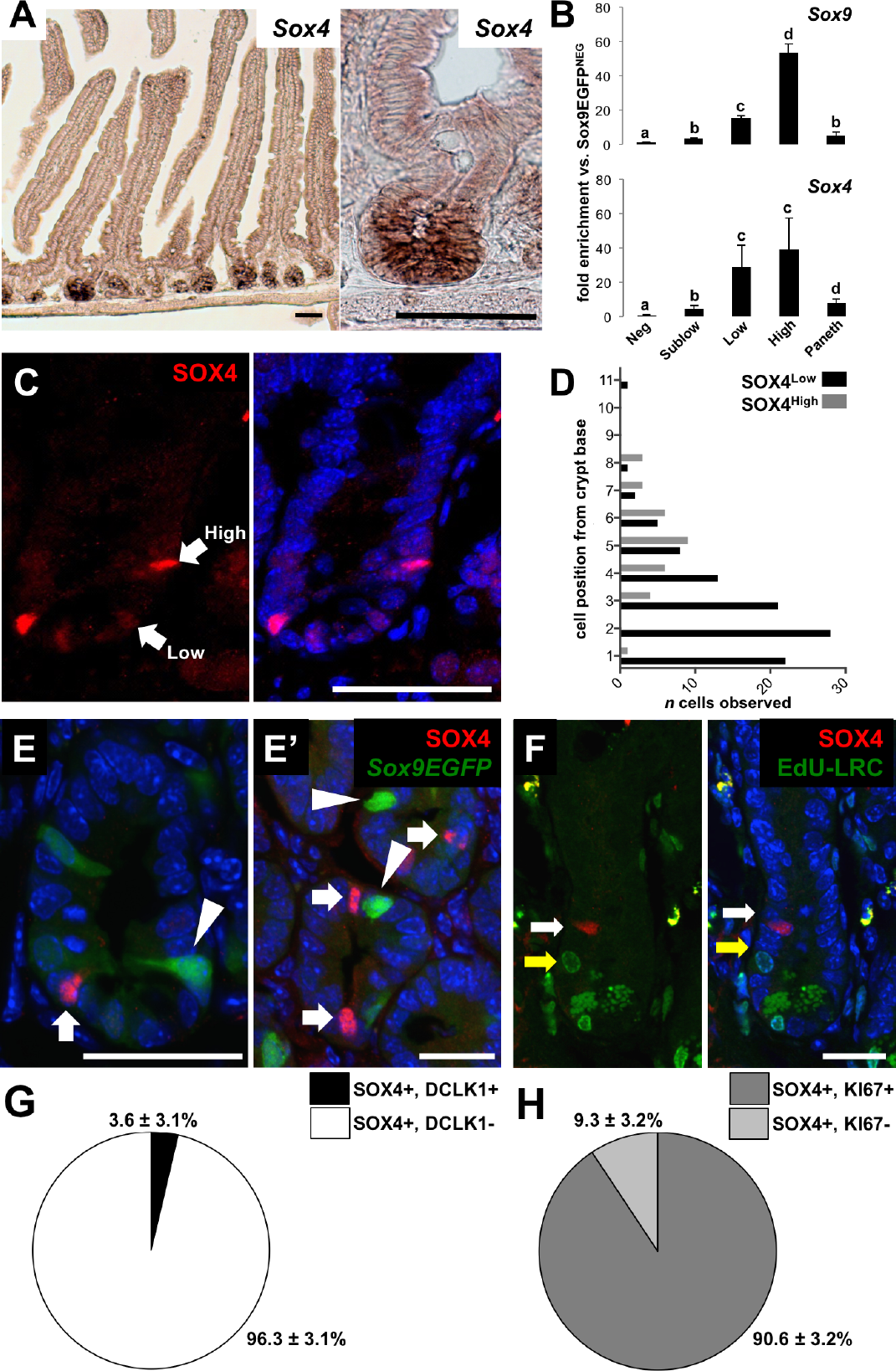
Sox4 is expressed in the ISC and early progenitor zone. (A) *In situ* hybridization localizes *Sox4* mRNA to the base of small intestinal crypts. (B) FACS isolation of *Sox9*^*EGFP*^ populations demonstrates enrichment of *Sox4* in active ISCs (*Sox9*^*low*^) and secretory progenitors/reserve ISCs (*Sox9*^*high*^) (different letters indicate statistically significant differences, p < 0.05). (C) SOX4 protein is expressed in the ISC and supra-Paneth cell zones at variable levels, with (D) SOX4^low^ cells primarily localized to the CBC position (1-3) and SOX4^high^ cells occurring mainly in the supra-Paneth cell zone (4+). (E/E’) SOX4^high^ and *Sox9*^*high*^ cells do not co-localize; arrows indicate SOX4hi cells and arrowheads indicate *Sox9*^*high*^ cells (n=3 mice, 100 SOX4^high^ cells per mouse, scale bar represents 20um). (F) SOX4^high^ cells are not LRCs by 28d continuous EdU labeling followed by 10d washout; white arrow indicates SOX4^high^ cell and yellow arrow indicates LRC (scale bar represents 20um; note: EdU detection kit reacts with Paneth cell granules). (G) A small portion of SOX4^high^ cells co-localize with the tuft cell marker DCLK1 and (H) a majority of SOX4^high^ cells are positive for proliferative marker KI-67.

Next, we examined SOX4 protein expression by immunofluorescence. SOX4 protein was expressed in a more restricted pattern relative to mRNA. Like *Sox9*, we observed distinct expression levels of SOX4, which we characterized as “low” and “high” (Figure 1C). SOX4^low^ cells were associated with the crypt base columnar (CBC), active ISC population, in cell positions +1-3 counting from the base of the crypt (Figure 1C, D). Interestingly, SOX4^high^ cells were predominantly found in the supra-Paneth cell position, which is commonly associated with early secretory progenitors, at cell positions +4-7 (Figure 1C, D) ^25^. Unlike SOX9, which is expressed in villus-based EECs/tuft cells, we did not detect any *Sox4* mRNA or protein outside of the crypts ^13^.

Since *Sox9*^*high*^ cells are known to represent EEC-like secretory progenitors, we asked if the SOX4^high^ population overlapped with *Sox9*^*high*^ cells. However, we found no co-localization between high levels of SOX4 protein and high levels of *Sox9* mRNA, as determined by *Sox9*^*EGFP*^, which is an accurate surrogate of endogenous *Sox9* mRNA and protein expression (*n* = 3 mice, 100 SOX4+ cells per mouse) (Figure 1E, E’) ^13, 26^. *Sox9*^*high*^-expressing cells are consistent with a subset of label-retaining reserve ISCs (LRCs) located in the supra-Paneth cell position. However, not all LRCs are *Sox9*^*high*^ ^23^. We reasoned that the supra-Paneth cell location of SOX4^high^ cells suggests that they may represent the non-*Sox9*^*high*^ subset of LRCs. To examine this, we counted the number of SOX4^high^ cells co-localizing with LRCs, as determined by 28 days of continuous labeling with 5-ethynyl-2-deoxyuridine (EdU), followed by 10 days of washout ^23, 25^. Surprisingly, we found that no SOX4^high^ cells were LRCs (*n* = 3 mice, 100 SOX4+ cells per mouse) (Figure 1F). We observed rare co-localization between DCKL1, a tuft cell/reserve ISC biomarker also associated with the +4 position, and SOX4 (5.0% ± 3.0%; Figure 1G). Finally, we addressed the possibility that SOX4^high^ cells may represent a quiescent population by co-staining with KI67, which marks all proliferating cells. 90.6 ± 3.2% of SOX4^high^ cells were found to be proliferative (Figure 1H). Together, these data suggest that SOX4^high^ cells represent ISCs or progenitors, and are distinct from LRC *Sox9*^*high*^ secretory precursors. Co-localization with DCLK1 suggests that a subset of SOX4^high^ cells may represent tuft cell or non-LRC secretory precursors.

### Loss of Sox4 results in increased proliferation and decreased ISC function

Complete deletion of *Sox4* results in embryonic lethality ^27^. To enable examination of *Sox4* function in adult intestinal epithelium, we crossed previously generated *Sox4^fl/fl^* mice to mice expressing Cre recombinase driven by the *villin* promoter (*vilCre*), which is constitutively expressed in the intestinal epithelium beginning at E13.5 ^28, 29^. To examine *Sox4* expression, we isolated CD44- and CD44+ populations from control and Sox4cKO adult intestinal epithelium (CD326+) by FACS and confirmed CD44 enrichment by qPCR (Supplementary Figure 1A-C). CD44 marks crypt cells as well as EECs in the villi (Supplementary Figure 1A). As expected, control animals exhibited significant enrichment of *Sox4* in the CD44+ population, consistent with restricted expression of *Sox4* in the intestinal crypts (Supplementary Figure 2A). Sox4cKO animals did not express detectable levels of *Sox4* mRNA or protein (Supplementary Figure 2A, B).

**Figure 2.**
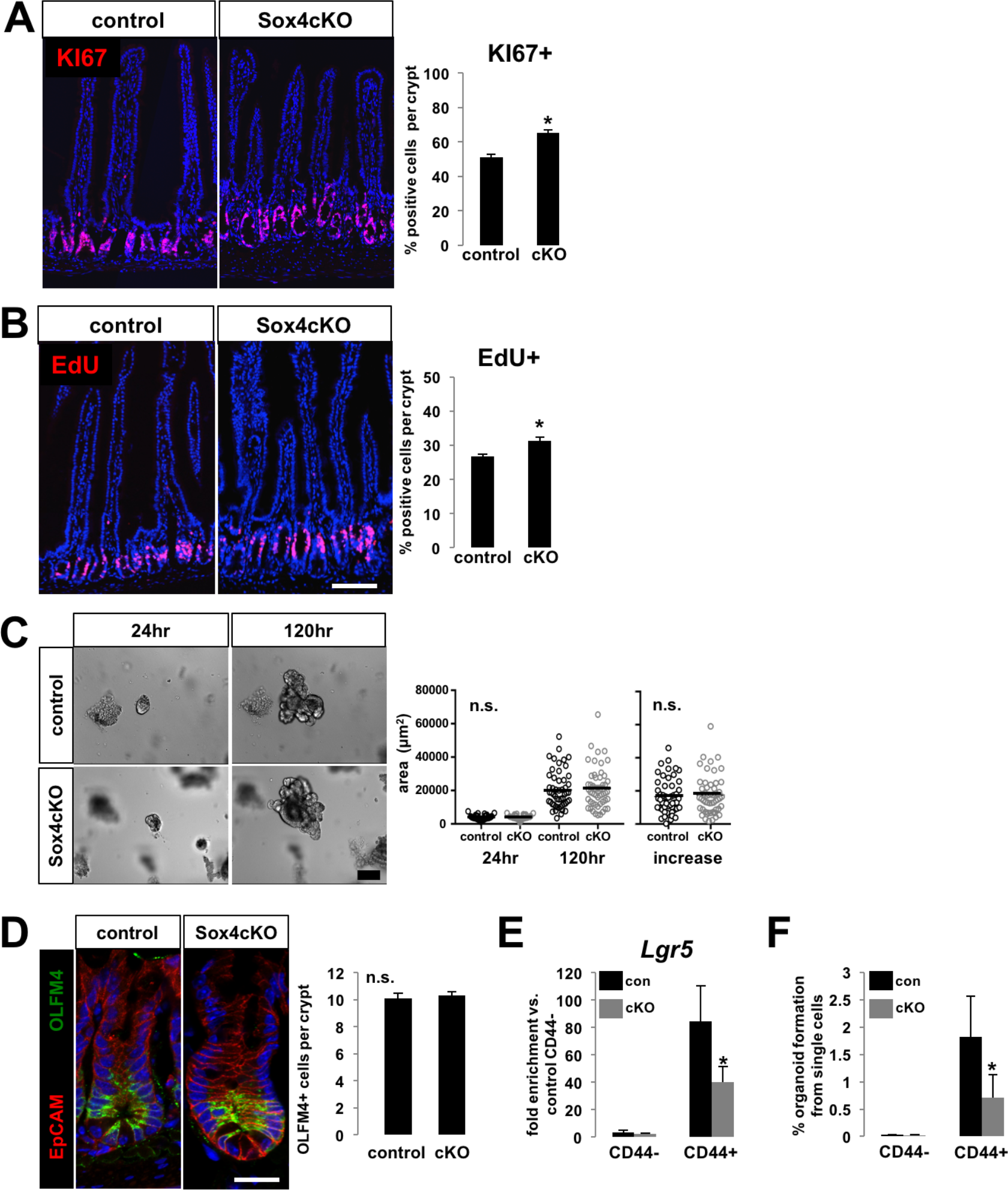
Loss of Sox4 results in increased proliferation and decrease in functional stemness, with no change in ISC numbers. (A) Sox4cKO intestines exhibit increased total proliferation by KI67 staining, as well as (B) increased numbers of cells in S-phase, as indicated by EdU (scale bar represents 100um). (C) Whole crypts isolated from Sox4cKO intestines form organoids that undergo similar growth rates to control crypts (scale bar represents 100um). (D) Numbers of OLFM4+ ISCs remain unchanged in Sox4cKO crypts, while (E) *Lgr5* is downregulated in CD44+ populations from Sox4cKO intestines. Functional stemness, as assayed by organoid forming ability, is significantly decreased in CD44+ populations isolated from Sox4cKO intestines (scale bar represents 25um; asterisks indicate significance; p < 0.05).

To assay the role of *Sox4* in ISC/TA proliferation, we examined KI67 and EdU incorporation, which was administered as a single, 2hr pulse. Sox4cKO crypts exhibited significantly higher percentages of total proliferating cells as well as cells in S-phase (Figure 2A, B). This increase in proliferation was reflected by increased crypt length in Sox4cKO intestines (Supplementary Figure 2C). In organoid cultures, Sox4cKO crypts grew at the same rate as control crypts, suggesting that hyperproliferation in the absence of *Sox4* may be masked by exogenous growth factors required for organoid culture (Figure 2C).

Due to its known role in maintenance of stem/progenitor pools in other tissues, we sought to determine if *Sox4* regulates the size or function of the ISC pool. OLFM4, which is an established marker of ISCs, was detected by immunofluorescence to assess ISC numbers, which were unchanged between control and Sox4cKO intestines (Figure 2D) ^30^. Next, we FACS-isolated CD44- and CD44+ cells from control and Sox4cKO mice and conducted single cell organoid forming assays to test ISC function ^31^. As expected, CD44-cells failed to form organoids regardless of *Sox4* status (Figure 2E). CD44+ cells from Sox4cKO mice produced approximately 50% fewer organoids than controls, demonstrating a significant loss of stemness (Figure 2E).

To determine if this functional deficit was coincident with an increase in apoptosis in Sox4cKO samples, we examined cleaved-caspase-3 expression in the crypts of control and Sox4cKO intestines, as well as the number of apoptotic cells in CD44- and CD44+ populations of cells prepared for FACS. In both cases, we found no significant difference in apoptosis between control and Sox4cKO samples (Supplementary Figure 3A, B). However, CD44+ cells from Sox4cKO mice expressed significantly lower levels of *Lgr5*, consistent with decreased stem cell function (Figure 2F). Together, these data indicate that *Sox4* does not regulate ISC numbers or apoptosis, but is required for normal ISC function and *Lgr5* expression in the intestinal crypt.

**Figure 3.**
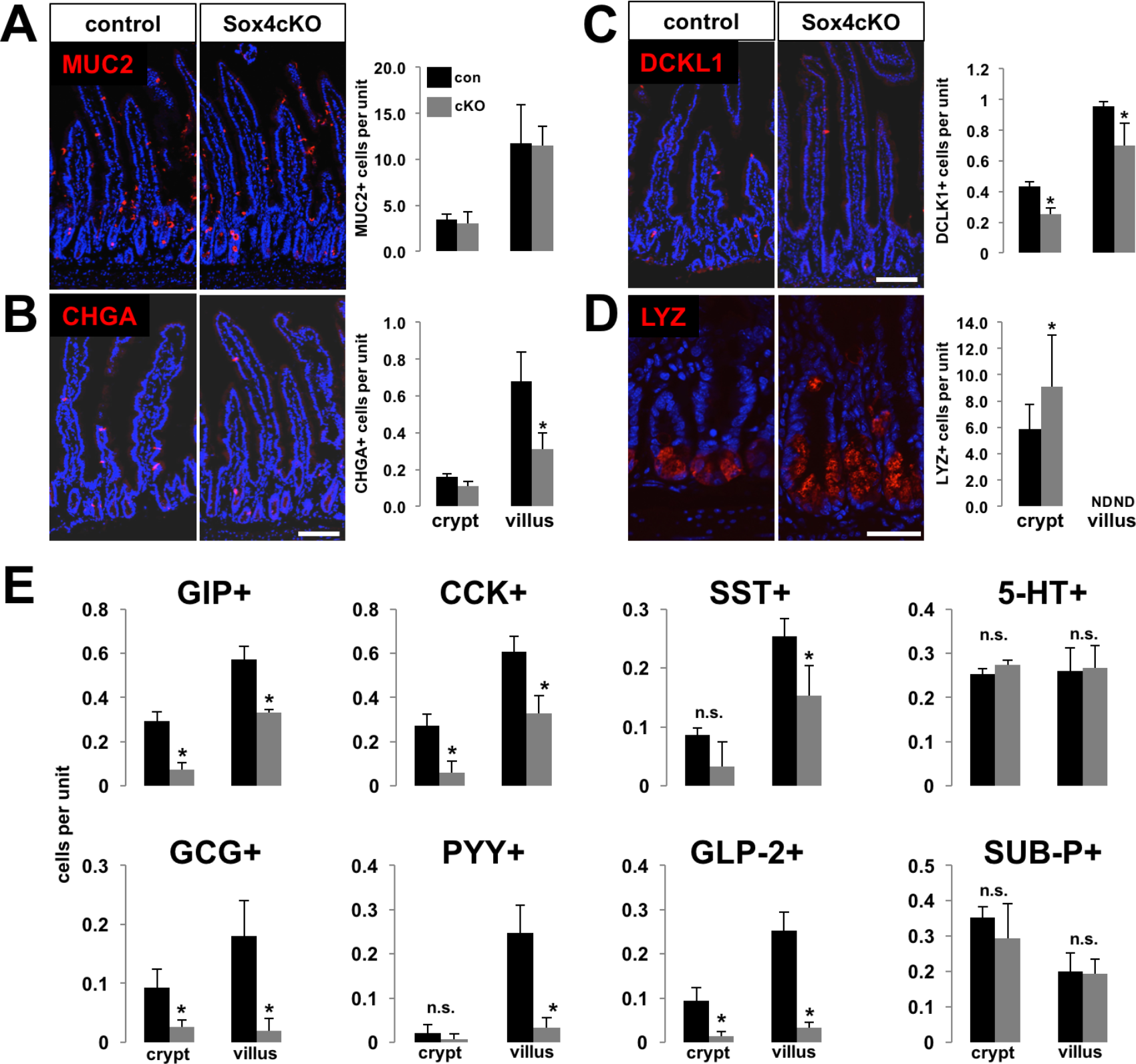
Sox4 is required for proper secretory lineage allocation. (A) MUC2+ goblet cells remain unchanged in Sox4cKO intestines, while (B) CHGA+ EECs are found at decreased numbers in Sox4cKO villi, and (C) DCLK1+ tuft cells are decreased in both crypt and villus compartments of Sox4cKO intestines (scale bar represents 100um). (D) Paneth cell numbers are increased in Sox4cKO crypts (scale bar represents 50um). (E) Assessment of EEC differentiation by immunofluorescence against markers associated with specific EEC subtypes reveals that *Sox4* is required for proper specification of a subset of EEC lineages. Conversely, numbers of cells positive for serotonin (5-HT) and Substance P (SUB-P) are unaffected in Sox4cKO intestines. (asterisks indicate significance; p < 0.05).

### Sox4 regulates secretory lineage allocation

We next sought to determine if *Sox4* regulates differentiation and intestinal lineage allocation. Goblet cells, detected by expression of MUC2, were found at similar numbers between Sox4cKO and control intestines (Figure 3A). CHGA^+^ EECs were found in significantly reduced numbers in the villi of Sox4cKO mice (Figure 3B). Numbers of DCLK1^+^ tuft cells were significantly reduced in crypt and villus compartments of Sox4cKO intestines (Figure 3C). Paneth cells were increased in Sox4cKO intestines (Figure 3D), reflecting additional Paneth cells at higher positions in Sox4cKO crypts (Supplementary Figure 4). Staining patterns for sucrose isomaltase (SIS), a brush-border enzyme associated with absorptive enterocytes, were unchanged between control and Sox4cKO samples (Supplementary Figure 5A). Additionally, villus height was unchanged, and no change was observed for *Hes1*, which specifies absorptive enterocyte fate (Supplementary Figure 2C and Supplementary Figure 5B).

**Figure 5.**
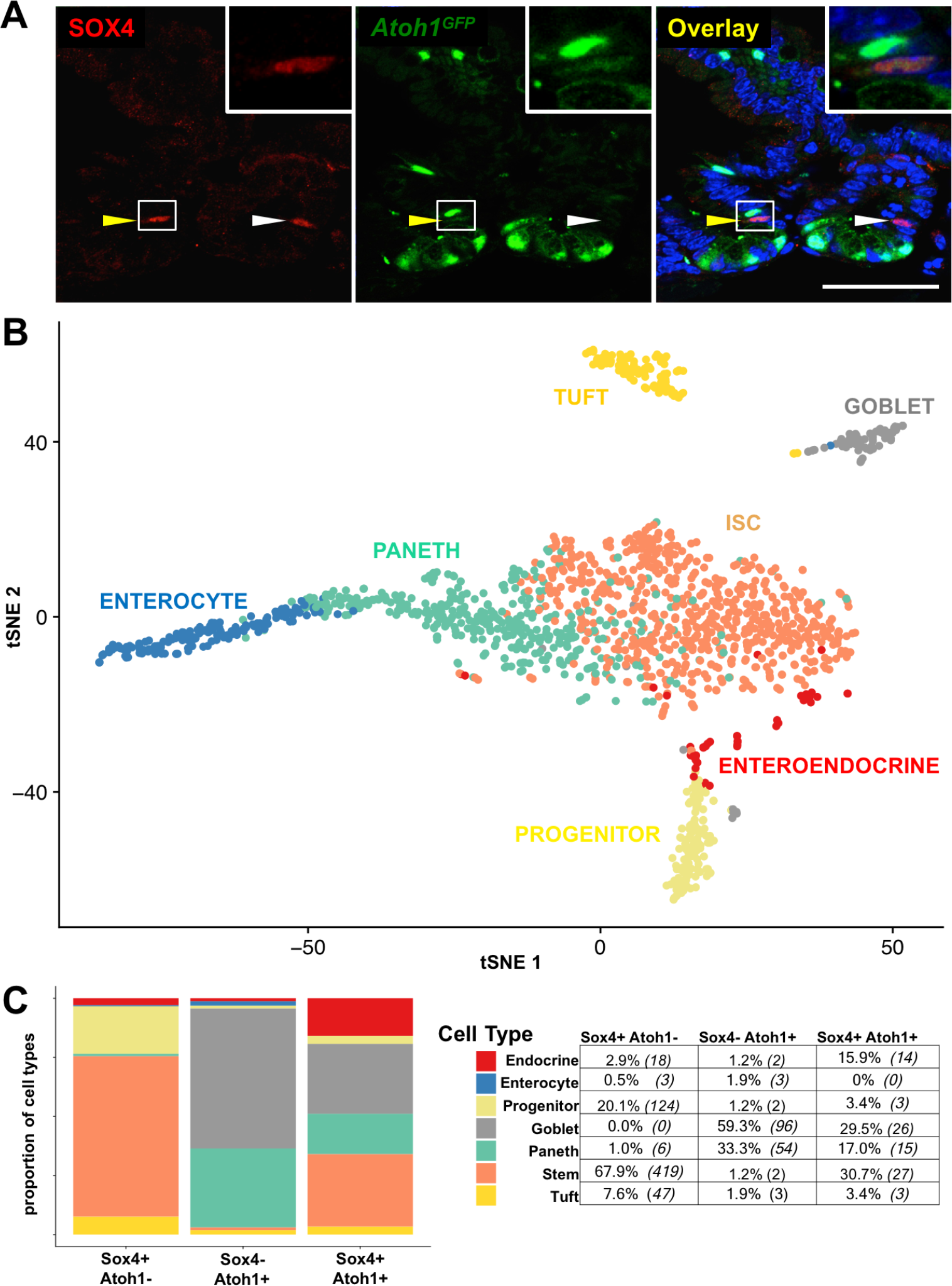
Atoh1+ and Sox4+ exhibit overlapping and exclusive expression in secretory cells and ISCs. (A) Co-detection of SOX4 and *Atoh1*, using a transgenic *Atoh1*^*GFP*^ allele, reveals that 55.3 ± 6.1% of SOX4+ cells co-express *Atoh1* (yellow arrowhead indicates SOX4+/Atoh1+ cell; white arrowhead indicates SOX4+/Atoh1-cell. Scale bar represents 25um). (B) scRNA-seq ^37^ identifies unique cell populations in the intestinal epithelium (C) Classifying cells based on *Sox4* and *Atoh1* expression status (*Sox4+*:*Atoh1-*, *Sox4-*: *Atoh1+*, and *Sox4+*:*Atoh1*+) reveals that *Sox4+* populations contain the highest proportion of tuft cells, EECs, and ISCs, consistent with observed phenotypes *in vivo* and *in vitro*.

In order to examine gene regulatory changes in Sox4cKO samples in an unbiased manner, we subjected FACS isolated CD44^-^ and CD44^+^ populations to RNA-seq. Consistent with histological observations, decreased expression of genes associated with EEC subpopulations, including pro-EEC transcription factors *Isl1, Nkx2.2,* and *Pax6*, was the predominant change observed in Sox4cKO CD44^+^ populations (Supplementary Figure 1D and Supplementary Table 1) ^32, 33^. There were no significant gene expression changes in the CD44^-^ populations consistent with the restricted expression pattern of SOX4 in CD44^+^ intestinal crypts and CD44+ status of EECs (Supplementary Figure 1D). Since the EEC lineage is comprised of a diverse complement of subpopulations, we re-examined Sox4cKO intestines for the presence of hormones produced by specific EEC subtypes. Compared to the modest reduction in CHGA^+^ cell numbers, we found substantial and significant decreases in EEC subpopulations expressing GIP, CCK, SST, GCG, PYY, and GLP-2 (Figure 3E). Interestingly, numbers of 5-HT and SUBP-expressing populations remained unchanged (Figure 3E).

Since Notch repression is required for EEC specification, we asked if *Sox4* is activated following downregulation of Notch. To do so, we treated wild type organoids with the gamma secretase inhibitor DAPT. Since *Sox4* has been reported to be regulated by Wnt in colon cancer cell lines and inhibited by Bmp in hair follicle progenitors, we also treated organoids with WNT3A and Noggin, a Bmp antagonist ^34, 35^. While *Sox4* levels and cell numbers did not respond to Wnt activation or Bmp inhibition, we found that Notch inhibition drove a 2-fold increase in *Sox4* expression that coincided with decreased expression of *Ascl2*, and ISC marker, and increased expression of *Ngn3*, and early EEC transcription factor (Supplementary Figure 6). Together, these data demonstrate that *Sox4* is required for proper secretory lineage allocation. Furthermore, *Sox4* is upregulated following Notch inhibition, consistent with early secretory differentiation responses, and required for proper specification of EEC subpopulations.

### Sox4 contributes to tuft cell differentiation and hyperplasia

Since a subset of SOX4^high^ cells were found to co-localize with DCLK1 and DCLK1^+^ tuft cells were decreased in Sox4cKO intestines, we sought to examine the regulatory role of *Sox4* in tuft cell differentiation more closely. IL-13 is known to induce tuft cell differentiation, and is produced by Group 2 innate lymphoid cells (ILC2) that are recruited to the intestinal epithelium in response to tuft cell secreted IL-25. This mechanism produces an extrinsic feed-forward loop that signals through unknown intrinsic response elements in ISCs and progenitors to drive tuft cell hyperplasia following parasitic helminth infection ^2, 5^. We treated wild type intestinal organoids with IL-13 to determine its effect on *Sox4*. As previously reported, IL-13 induced expression of tuft cell-specific *Dclk1* (Supplementary Figure 7A). Compellingly, IL-13 also induced a dose-dependent upregulation of *Sox4*, driving an approximately 3-fold increase at the optimal dose (Supplementary Figure 7A). Levels of *Sox4* upregulation correlated closely with *Dclk1* induction. As expected, goblet cell *Muc2* was the only non-tuft lineage specific gene also upregulated in response to IL-13 (Supplementary Figure 7B) ^2^. To determine if *Sox4* is required for proper *Dclk1* upregulation in response to IL-13, we next treated Sox4cKO and control organoids with IL-13. While control organoids exhibited robust induction of *Dclk1* and the tuft cell transcription factor *Pou2f3*, Sox4cKO organoids failed to significantly upregulate these genes (Figure 4A).

**Figure 4.**
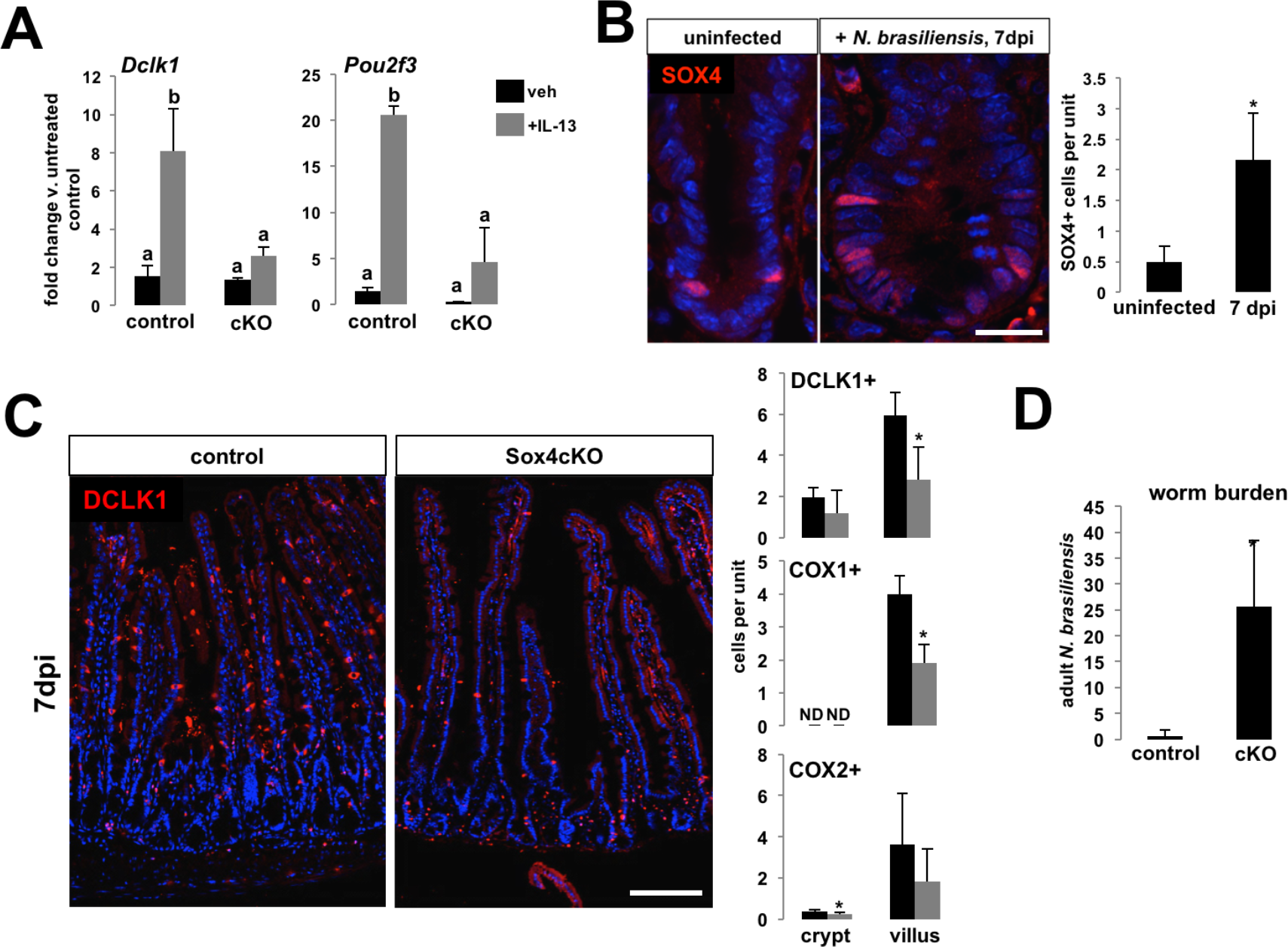
Sox4 regulates tuft cell differentiation and is required for tuft cell hyperplasia following helminth infection. (A) Tuft cell markers *Dclk1* and *Pou2f3* are upregulated in control, but not Sox4cKO organoids, in response to treatment with recombinant IL-13 (different letters indicate statistically significant differences, p < 0.05). (B) 7 days post-infection with *N. brasiliensis*, control intestinal crypts exhibit a significant expansion of SOX4^high^ cells (scale bar represents 20um). (C) Control mice demonstrate expected tuft cell hyperplasia at 7dpi, where Sox4cKO intestines have significantly fewer tuft cells as quantified by DCLK1, COX1, and COX2 (ND = no data; COX1+ cells are not found in intestinal crypts. Scale bar represents 100um). (D) Sox4cKO mice also fail to fully clear adult worms by 7dpi (asterisks indicate significance, p < 0.05, n=3 mice per group).

Next, we asked if *Sox4*-driven tuft cell responses were physiologically necessary for clearance of helminth infection. Sox4cKO and control animals were infected with *N. brasiliensis*, a helminth known to induce tuft cell-mediated type II immune reactions ^2, 4, 5^. Intestines were examined 7 days post-infection (d.p.i.), at the peak of tuft cell hyperplasia, when most worms are typically cleared from wild type mice ^5^. We first examined SOX4^+^ cells in control mice. Helminth infection drove a 4-fold increase in the number of SOX4^+^ cells per crypt (Figure 4B). As expected, DCLK1^+^ cell numbers were largely upregulated in control animals (Figure 4C). In contrast, Sox4cKO intestines exhibited an attenuated increase in tuft cell numbers (Figure 4C). To determine if impaired tuft cell hyperplasia in Sox4cKO animals affected worm burden at 7d.p.i., we quantified the number of adult worms present in the intestines of infected animals. In order to match worm clearance data with tuft cell hyperplasia in the same mouse, duodenal and ileal tissues were harvested to establish worm counts and jejunal tissues were used for histological studies, discussed above. Control animals exhibited a near-total clearance of adult worms, while Sox4cKO animals bore a significantly higher worm burden at 7d.p.i. (Figure 4D). These data implicate *Sox4* as a regulator of tuft cell differentiation at physiological baseline, with an important role in tuft cell hyperplasia during helminth infection.

### Sox4- and Atoh1-expressing cells form overlapping and distinct populations

Early tuft cell specification remains a paradox in intestinal epithelial differentiation. *Atoh1* is accepted as the master regulator of early secretory fate in the intestine, and is de-repressed following loss of Notch signaling ^36^. While tuft cells fail to differentiate properly in the presence of constitutively active Notch, multiple lines of evidence demonstrate that a majority of tuft cells are not derived from *Atoh1*-expressing progenitors ^3, 6^. These data suggest the existence of Notch-repressed, *Atoh1*-independent tuft cell regulatory pathways that remain uncharacterized. Since our data demonstrate that *Sox4* is upregulated following Notch inhibition and is involved in tuft cell differentiation, we hypothesized that it might be expressed in an *Atoh1*-independent manner. We examined SOX4 expression in *Atoh1*^*GFP*^ intestines, and found that 55.3 ± 6.1% of SOX4^high^ cells were *Atoh1*-negative, with the remainder co-localizing with Atoh1-GFP (Figure 5A).

To examine *Sox4* and *Atoh1* populations more thoroughly, we reanalyzed previously published single cell RNA-seq (scRNA-seq) data from primary mouse intestinal epithelial cells ^37^. Cells were visualized by tSNE dimensionality reduction and classified into previously characterized phenotypic populations (Figure 5B) ^37^. Next, we queried the distribution of cell types within each of three populations of interest: (1) *Sox4+*:*Atoh1*-(*n* = 617 cells), (2) *Sox4+*:*Atoh1*+ (*n* = 88 cells), and (3) *Sox4-*:*Atoh1*+ (*n* = 162 cell). Within the *Sox4*+ population, 12.5% of cells were also *Atoh1*+. Next, we determined the cellular composition of each of our populations of interest. The *Sox4+*:*Atoh1*-population consisted primarily of ISCs and progenitor cells (88%), but also contained the highest proportion of tuft cells relative to other populations (Figure 5C). The *Sox4+*:*Atoh1*+ population was a mix of ISCs and secretory cell types, and contained the highest proportion of EECs relative to other populations (Figure 5C). As expected, the *Sox4-*:*Atoh1*+ population was comprised mostly of secretory cells, with the lowest proportion of ISCs an progenitor cells and the highest proportion of goblet and Paneth cells relative to other populations (Figure 5C). These data demonstrate that *Sox4* and *Atoh1* form distinct and overlapping populations, and that *Sox4* is most strongly correlated with ISCs, progenitors, and tuft cells when expressed exclusively of *Atoh1* and more strongly correlated with EECs when co-expressed with *Atoh1*.

### Sox4 drives tuft cell differentiation independently of Atoh1

We next sought to determine if *Sox4* is sufficient to drive secretory EEC and tuft cell differentiation. To this end, we generated stable transgenic organoids carrying a cumate-inducible *Sox4* overexpression vector (Sox4OE). Gain of function was validated by immunofluorescent detection of SOX4 in empty vector control and induced Sox4OE organoids (Supplementary Figure 8). To validate organoid models for assessment of *Sox4*-dependent differentiation phenotypes, we included organoids isolated from Sox4cKO mice in our analyses. As expected, Sox4cKO organoids exhibited very low numbers of DCLK1+ tuft cells and CHGA+ EECs (Figure 6A, B). In contrast, overexpression of *Sox4* resulted in a significant increase in numbers of both tuft cells and EECs, relative to empty vector controls (Fig 6A, B).

**Figure 6.**
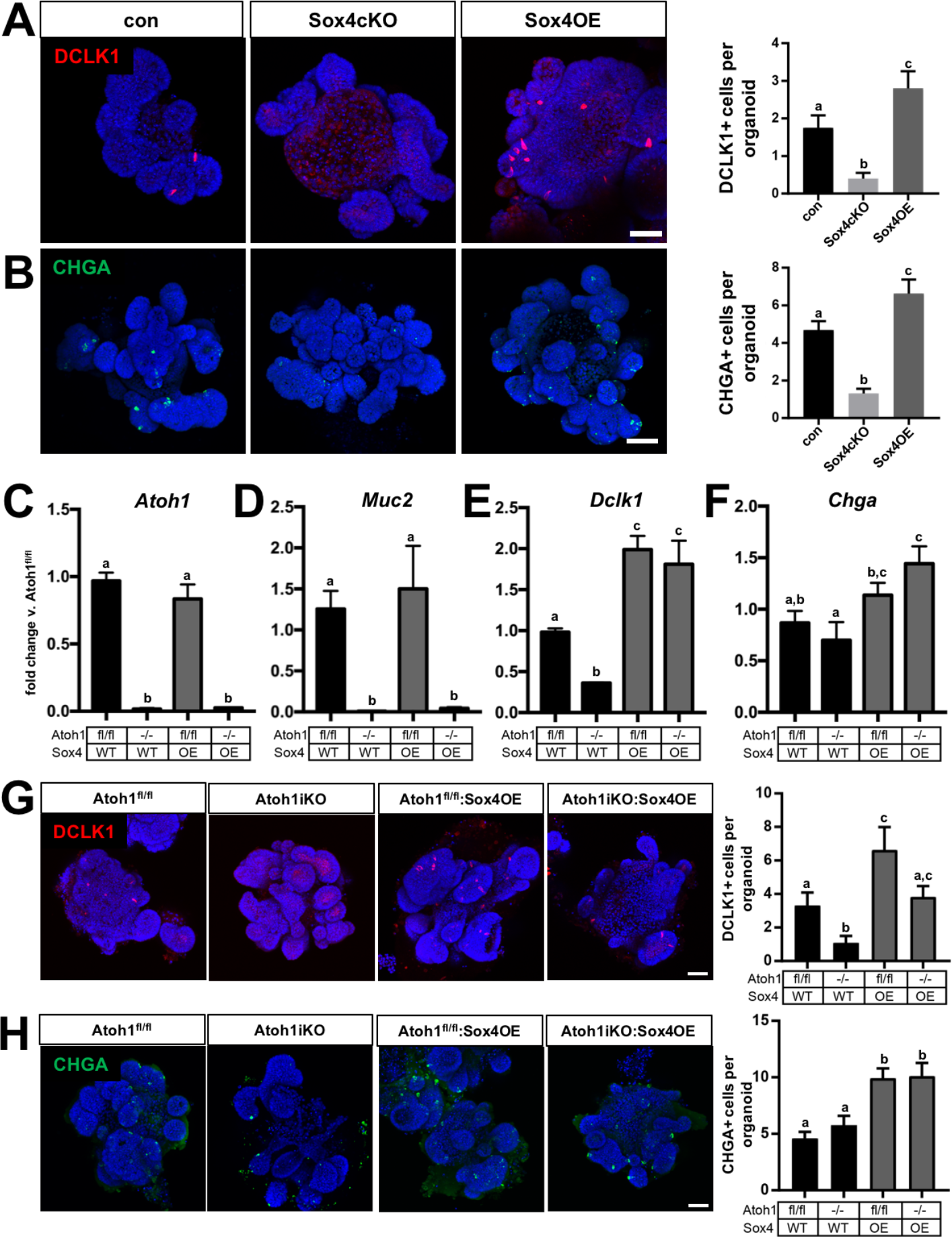
Sox4 drives tuft cell and EEC differentiation independently of Atoh1. (A) Tuft cells, as identified by DCLK1, are rare in control organoids, absent in Sox4cKO organoids, and upregulated in Sox4OE organoids. (B) Similar results are observed for CHGA+ EECs, indicating that *Sox4* is sufficient to increase tuft and EEC differentiation. 4-OHT treatment downregulates *Atoh1* in control and Sox4OE organoids, resulting in loss of *Muc2* expression, which demonstrates loss of *Atoh1*-driven transcription. (E) Loss of *Atoh1* in organoids also results in downregulation of *Dclk1*, which is rescued to normal levels in Sox4OE organoids, regardless of *Atoh1* expression. (F) *Chga* is not affected by *Atoh1* deletion in organoids, but is upregulated by Sox4OE in the absence of *Atoh1*. Cell counts of (G) DCLK1+ tuft cells and (H) CHGA+ EECs demonstrate correlation between changes in gene expression and lineage specification in Atoh1iKO and Sox4OE organoids. Notably, CHGA+ cells numbers are increased in Sox4OE organoids both with and without *Atoh1* expression.

Since scRNA-seq demonstrated differential distribution of *Sox4* and *Atoh1* across progenitors and secretory cell types, we hypothesized that *Sox4* may function independently of *Atoh1* in secretory differentiation. To test this, we generated Sox4OE organoids derived from Atoh1^fl/fl^:vilCre^ER^ mice. *Atoh1* recombination was induced by treating organoids with 1uM 4-OHT once a day for three days and collecting for analysis 24hr following final treatment with 4-OHT. Analysis of *Atoh1* expression confirmed the effectiveness of this strategy in generating Atoh1iKO organoids (Figure 6C). To assess whether residual ATOH1 protein was exerting continued effects in Atoh1iKO organoids, we next analyzed *Muc2* expression, which is associated with Atoh1-dependent goblet cell differentiation. We found that *Muc2* levels were also significantly downregulated in Atoh1iKO organoids, suggesting no residual ATOH1 function at the time of organoid analysis (Figure 6D).

Next, we examined the effect of *Atoh1* and *Sox4* on tuft and EEC differentiation. Surprisingly, *Dclk1* was significantly downregulated following loss of *Atoh1*, in direct conflict with reported *in vivo* data demonstrating an increase in tuft cells in *Atoh1*-null intestines (Figure 6E) ^10^. However, treatment of Atoh1iKO organoid cultures with IL13 restored *Dclk1* expression to control levels, in contrast to the loss of IL13 responsiveness in Sox4cKO organoids (Supplementary Figure 9). Overexpression of *Sox4* was also sufficient to restore *Dclk1* expression levels in the absence of *Atoh1* (Figure 6E). Unexpectedly, *Chga* expression levels remained unchanged following loss of *Atoh1*, and were upregulated by *Sox4* in Atoh1iKO organoids (Figure 6F). Notably, Sox4OE failed to rescue or upregulate *Muc2* expression levels, suggesting specificity to tuft and EEC lineages (Figure 6D). To refine these observations, we conducted whole mount staining on Atoh1iKO and Sox4OE organoids and directly quantified tuft and EEC numbers in whole organoids. Quantification by immunofluorescence recapitulated gene expression results, demonstrating upregulation of DCLK1+ tuft cells and CHGA+ EECs in Sox4OE organoids regardless of *Atoh1* expression (Figure 6G and H). Together, these data demonstrate that *Sox4* is sufficient to drive an increase in tuft and EEC differentiation and can do so independently of *Atoh1*.

## Discussion

Here we show that genetic ablation of *Sox4* in the intestinal epithelium results in increased numbers of proliferating cells and Paneth cells, as well as a reduction in EEC and tuft cell lineages. Our study supports *Sox4* mediated regulation of stem cells in other organ systems by demonstrating that *Sox4* regulates ISC function and differentiation toward EECs and tuft cells in the intestinal epithelium. Additionally, we demonstrate that *Sox4* has an essential role in pushing lineage allocation toward tuft cell differentiation in response to helminth infection, implying an important role for *Sox4* in host responses to pathogens.

Our group and others have reported *Sox4* expression in *Lgr5+* CBCs, but a role for *Sox4* in ISCs has yet to be established ^18, 22^. We demonstrate that the numbers of ISCs quantified by OLFM4 immunostaining are unchanged between Sox4cKO and control samples. These data suggest that the associated increase in proliferating cells in Sox4cKO crypts is likely due to increased progenitor numbers and not more robust ISC activity. Rather, *in vitro* assay of ISC function demonstrated impaired organoid-forming efficiency of single ISCs isolated from Sox4cKO mice. As this dysfunction is not associated with a change in the number of apoptotic cells between control and Sox4cKO mice, it is likely that decreased organoid-forming efficiency is caused by loss of stemness and premature differentiation in *Sox4* deficient ISCs. In this regard, Sox4cKO mice demonstrate reduced *Lgr5* mRNA. *Lgr5* is highly expressed in ISCs and serves as a receptor for R-Spondin ligands that potentiate Wnt-signaling, which is in turn required to maintain stemness ^38^. These data indicate that *Sox4* is not required for maintaining ISC numbers, but suggests that *Sox4* plays a role in maintaining stemness by positively regulating *Lgr5* levels. Functionally, this role would be consistent with *Sox4* knockout phenotypes in other tissues, including the developing heart, skeletal muscle, nervous, renal, and hematopoietic systems, which often exhibit precocious differentiation and reduced progenitor and post-mitotic cell numbers ^19, 20, 27, 39-41^.

Sox4cKO intestines also demonstrate decreased EEC and tuft cell numbers, implying that *Sox4* plays a key role in the specification of these secretory lineages. RNA-seq analysis demonstrates that *Sox4* regulates the expression of the EEC transcription factors, including *Isl1, Pax6,* and *Nkx2-2*. *Isl1* deletion in the intestine is known to lead to loss of EECs subtypes expressing GIP, CCK, SST, GCG, PYY, and GLP-2 ^33^. We demonstrate that loss of *Sox4* results in a marked reduction of *Isl1* expression with a concomitant reduction in the numbers of the same EEC subtypes. Together, these results indicate that *Sox4* likely functions upstream of EEC transcription factors *Isl1, Pax6*, and *Nkx2-2* to drive EEC specification.

The mechanisms that give rise to the tuft cell lineage are largely understudied due to the relatively new characterization of tuft cells as a distinct lineage. The role of intestinal tuft cells was recently described by a handful of independent studies, all of which demonstrated tuft cell hyperplasia following infection with parasitic helminths.

Following infection, tuft cells initiate a type II immune response through ILC-2 cells ^2, 4, 5^. ILC-2 cells secrete IL-13, which induces tuft cell hyperplasia ^2, 4, 5^. *Pou2f3* is known to be essential for tuft cell differentiation, but is also expressed by mature, villus-based tuft cells ^2^. Other transcription factors involved in tuft cell specification and hyperplasia remain unknown, and tuft cell progenitors have no distinct biomarkers ^2^.

Though SOX4^high^ cells are relatively rare in homeostasis, we observed a striking increase in SOX4^high^ cell numbers coincident with peak tuft cell hyperplasia following parasitic infection, suggesting that SOX4^high^ cells may represent a tuft progenitor population. We show that Sox4cKO epithelium demonstrates attenuated tuft cell hyperplasia, both in response to recombinant IL-13 *in vitro* and helminth infection *in vivo*, and that Sox4cKO mice are functionally impaired at clearing worms by 7d.p.i. Notably, IL13-mediated activation of *Pou2f3* is also impaired in Sox4cKO organoids, suggesting that *Sox4* acts upstream of this established tuft cell transcription factor. Since there is no reporter mouse model for *Sox4*, we leveraged existing scRNA-seq data to further interrogate the identity of *Sox4*+ cells. In agreement with histological data, we find that *Sox4* is enriched in ISC, progenitor, and secretory populations. Classifying *Sox4*+ cells by *Atoh1* expression reveals a strong correlation between *Sox4*+:*Atoh1*-cells and tuft cells and between *Sox4*+:*Atoh1*+ cells and EECs, supporting dual roles for *Sox4* in tuft and EEC differentiation. Functionally, this role is demonstrated by the ability of ectopic *Sox4* expression to induce increased tuft and EEC numbers in organoids.

Lineage specification of tuft cells from the ISC/progenitor pool remains a significant question in the field, as tuft cells are capable of being produced independently of *Atoh1*, but are inhibited by constitutive Notch signaling ^3, 6^. Further, it was recently shown that inducible knockout of *Atoh1 in vivo* leads to increased tuft cell numbers, implying that *Atoh1* may suppress tuft cell differentiation ^10^. Surprisingly, our *in vitro* analyses of Atoh1iKO organoids demonstrate the opposite effect. That is, loss of *Atoh1 in vitro* leads to decreased tuft cell numbers. However, in contrast to Sox4cKO organoids, which fail to upregulate *Dclk1* in response to exogenous IL13, Atoh1iKO organoids retain IL13-responsiveness in terms of *Dclk1* expression. These data suggest that tuft cells may be specified by multiple signaling events and pathways, but that IL13-mediated tuft cell hyperplasia is *Sox4*-dependent and *Atoh1*-independent. In agreement, our gain-of-function studies reveal that, although Atoh1iKO organoids exhibit reduced tuft cell numbers relative to non-recombined controls, *Atoh1* is dispensable for *Sox4*-driven tuft cell differentiation.

A surprising result of our studies is the observation that *Sox4* overexpression was sufficient to drive increased EEC differentiation in the absence of *Atoh1*. This effect does not appear to be the result of residual ATOH1 activity or non-specific *Sox4* regulation of all secretory differentiation, as goblet cell-associated *Muc2* expression remains severely downregulated in Atoh1iKO organoids at the timepoint examined, regardless of *Sox4* expression status. These data raise interesting questions regarding the role of *Atoh1* as a master regulator of secretory differentiation. To our knowledge, no other studies to date have expressed transcriptional regulators of secretory differentiation that are normally expressed upstream of *Atoh1* in the absence of *Atoh1*. Further studies in this area would be necessary to determine if other transcription factors are capable of rescuing secretory differentiation in Atoh1iKO models.

The existence of *Atoh1*-independent lineage specification of tuft cells and EECs challenges the long-standing dogma of *Atoh1* as the master transcriptional regulator of secretory differentiation in the intestine. Our findings demonstrate that *Sox4* is upregulated following downregulation of Notch or in response to IL13, and drives tuft or EEC specification in a manner that can proceed independently of *Atoh1*. These findings implicate *Sox4* as an important regulator of host responses parasitic infection and shed further light on epithelial components of Type II immune responses.

## Acknowledgments

The authors would like to thank Drs. Rashmi Chandra and Roger Liddle (Duke University, Durham, NC) for providing antibodies against CCK and PYY; Dr. Veronique Lefebvre for providing *Sox4^fl^* mice; Brian Golitz and Noah Sciaky for assistance with Illumina sequencing and analysis; Dr. Pablo Ariel and the UNC Microscopy Services Laboratory for assistance with confocal microscopy. RNA-seq data associated with this manuscript have been deposited in NCBI’s Gene Expression Omnibus and are available under GEO accession number GSE90795. This work was funded by the National Institutes of Health, R01 DK091427 (Magness), the Center for Gastrointestinal Biology and Disease P30 DK034987 (Magness), National Institutes of Health, R01 AI119004 (Reinhardt), and National Institutes of Health, R01 CA1428260, R01 DK092306 (Shroyer). A.D.G. was partially supported by the UNC GI Division Basic Science Training Grant, T32 DK007737-20 (Sartor), National Institutes of Health, K01 DK111709 (Gracz), and a Research Scholar Award from the American Gastroenterological Association (Gracz).

